# Genomic and Proteomic Analysis of Patients with Reoccurring Discrete Subaortic Stenosis

**DOI:** 10.1101/2023.01.22.525022

**Authors:** Ravi K. Birla, Kavya L. Singampalli, Asela Nieuwsma, Sunita Brimmer, Aditya Kaul, Lalita Wadhwa, Sung Y. Jung, Dereck Mezquita, Sandra Grimm, Swathi Balaji, Cristian Coarfa, Christopher Caldarone, Jane Grande-Allen, Sundeep G. Keswani

**Affiliations:** Laboratory for Regenerative Tissue Repair, Texas Children’s Hospital, Houston, Texas, USA; Center for Congenital Heart Surgery, Texas Children’s Hospital, Houston, Texas, USA; Department of Bioengineering, Rice University, Houston, Texas, USA; Medical Scientist Training Program, Baylor College of Medicine, Houston, Texas, USA; Division of Congenital Heart Surgery, Texas Children’s Hospital, Houston, Texas, USA; Department of Molecular and Cellular Biology, Baylor College of Medicine, Houston, TX, USA; Department of Biochemistry and Molecular Biology, Baylor College of Medicine, Houston, TX, USA; Department of Surgery, Baylor College of Medicine, Houston, Texas, USA; Division of Pediatric Surgery, Department of Surgery, Texas Children’s Hospital, Houston, Texas, USA

## Abstract

Discrete subaortic stenosis (DSS) is a pediatric condition in which a fibrotic membrane forms within the left ventricular outflow track. The fibrotic membrane is removed surgically; however, there is a high rate of reoccurrence which requires a second surgery. There are currently no tools available to predict the risk of reoccurrence in DSS patients, a limitation addressed by this study. In this study, we analyzed resected fibrotic membranes for DSS patients at the time of first surgery for non-recurrent and recurrent patients, and at the time of second surgery for recurrent patients. RNA-sequencing was conducted to obtain a global screen of changes in RNA expression while mass spectrometry was used to obtain a global screen of changes in protein expression. The results from the RNA-sequencing and mass spectrometry provide valuable insight into genes and the proteins that are differentially regulated in recurrent vs non-recurrent DSS patients.

## Introduction

Discrete sub-aortic stenosis (DSS) is a congenital heart condition in which a fibrous membrane forms in the left ventricular outflow track (LVOT) of patients ^1, 2^. This condition accounts for 6% of all congenital heart disorders and 8-30% of all cases of LVOT obstruction in the pediatric population ^3, 4^. The most significant clinical manifestation of DSS is an increase in peak LVOT pressure gradient and this in turn, can lead to chest pains, heart failure and syncope. The fibrous membrane is located just beneath the aortic valve, fusing with its leaflets and occasionally attached to the anterior mitral valve leaflet. DSS is associated with anatomical abnormalities in the LVOT, and one of the most common manifestations is a steep aortoseptal angle (AoAS) greater than 120°.

The acquired or inherited origin of DSS has not been established and continues to be a topic of debate, though there is greater evidence supporting the latter. The underlying inheritance genes linked to DSS have not been identified, therefore not supporting the inherited origin of DSS ^5, 6^. However, there is accumulating evidence supporting the acquired nature of DSS ^7, 8^, predominantly based on changes in hemodynamic conditions within the LVOT resulting from obstructed flow and subsequent remodeling of exposed endocardial endothelial cells and fibroblasts.

There are significant hemodynamic changes in DSS patients ^9^ resulting from LVOT obstruction due to the fibrotic membrane and steep AoAS ^10, 11^. Fluid structure interaction (FSI) models have been used to study the interaction of fluid shear stresses on the underlying fibrotic lesion and subsequent impact of these changes on the septal wall ^10^. Using FSI models, it has been shown that LVOT obstruction from an increased AoAS results in a 2.4-fold increase in peak velocity, 2.8-fold in vorticity, 5-fold increase in the magnitude of turbulent kinetic energy and a 23% increase in temporal shear magnitude of the septal wall ^10^. These changes demonstrate significant remodeling within the LVOT resulting from obstructed flow.

The molecular mechanisms responsible for the DSS are not known. However, anatomical abnormalities in the LVOT are hypothesized to initiate the formation of the fibrotic membrane while changes in the hemodynamics environment result in disease progression. While the cellular response to changes in LVOT anatomy and hemodynamics remain unknown, the role of mechanosensitive endocardial endothelial cells has been postulated. This hypothesis is based on the known response of endothelial cells to fluid shear stress ^12^, although a similar response has not been confirmed in DSS patients. The subsequent proliferative response of fibroblasts leading to the formation of the fibrous membrane has been established ^13^, though the trigger for this fibroproliferative response is not known.

The only treatment for DSS is surgical removal of the fibrotic lesion, considered to be a safe operation, although it is still associated with a 3% surgical mortality rate. The risks associated with surgery as those related to anesthesia, sternotomy and heart bypass. While initial surgery is a low-risk operation, there is a high rate of recurrent of the fibrotic membrane. Recurrent rates are 8-34% and require regular follow-up and constant monitoring of DSS patients. The risk factors for reoccurrence are young age at initial diagnosis and first surgery, higher preoperative peak LVOT gradient and sex, with females have a 1.5 times higher risk of reoccurrence compared with males^1^. The only way to treat patients with recurrent DSS is a second operation to remove the fibrotic membrane, associated with the same risks as the first operation.

Currently, there are no predictive tools to determine the risk of reoccurrence in DSS patients and as a result, all patients must be monitored aggressively. There is, however, one recent publication which correlates the rate of reoccurrence with known clinical metrics, like age at the time of first surgery, peak LVOT gradient and separation of the fibrotic membrane from the mitral valve and distance between fibrotic membrane and aortic valve ^14^. However, this study does little towards the development of predictive tools as it is based purely on empirical clinical data and the molecular response has not been factored. The ability to predict the probability of reoccurrence of the fibrotic lesion at the time of first surgery will dramatically impact patient care. For example, patients with a lower risk for reoccurrence will not require aggressive followup, and this will dramatically increase their quality of life.

In this study, we sought to develop a predictive tool for DSS reoccurrence. We sought to understand changes in the gene and protein levels for patient specific fibrotic lesions resected at the time of first and second surgery. Differential gene and protein expression was analyzed for DSS patients with different clinical outcomes. DSS patients with no reoccurrence 5 years after initial operation were termed *“non-aggressive DSS phenotype”*, while those with reoccurring phenotype within the same time frame were termed *“aggressive DSS phenotype”*. The fibrotic membranes from DSS patients with non-aggressive and aggressive phenotypes were processed for RNA-sequencing for genomic analysis and mass spectrometry for proteomic analysis. The objective of this study was to determine specific pathways, both at the gene and protein level, that are differentially expressed DSS patients with aggressive vs non-aggressive phenotype.

## MATERIALS AND METHODS

### Tissue Biopsies

The fibrous membrane is resected at the time of DSS operation, flash frozen and stored in −80°C in the TCH Biorepository according to IRB protocol H-26502. Samples were then transported to the lab on ice, divided into 4 equal pieces, one each for immunohistochemistry (IHC), RNA-sequencing, mass spectrometry for proteomics analysis and the 4^th^ specimen maintained as a reserve.

### Patient Population

For this study, we used tissue biopsies from 20 pediatric patients diagnosed with non-aggressive DSS phenotype and 12 patients with at least one recurrent phenotype requiring a second operation.

### ECHO Data

2D Doppler echocardiography was used to assess heart function and quantify changes in hemodynamics. The following parameters were obtained from the echocardiography data: heart rate, ejection fraction, left ventricle end diastolic volume (LVEDV), left ventricle end systolic volume (LVESV), LVOT peak gradient, LVOT mean gradient, aortic annulus, mitral valve annulus and aortaseptal angle.

#### RNA-Sequencing

total RNA was extracted from tissue biopsies using the RNeasy Micro Kit (Qiagen catalog # 74004), the purity was validated using the absorbance readings at 260 nm and 280 nm with 1.8-2.0 considered acceptable. Genomic analysis was conducted by the CPRIT Multi-omics Analysis Core, Advanced Technology Core at Baylor College of Medicine.

RNAseq data was trimmed using cutadap v1.18 and fastQC v0.11.9. Mapping was done with Homo_sapiens.GRCh38.101.gtf ^15^ as a reference genome. Trim and mapping quality was assessed with the multiqc ^16^ utility version 1.8. Differential expression analysis was done with use of the edgeR ^17^ package version 3.32.1 and EDAseq ^18^ 2.24.0. An FDR cutoff of 0.05 was selected and fold change cutoff: 1.5; LRT (likelihood ratio test) RUVr (remove unwanted variation) upperquartile normalization was used. GSEA ^19^ (gene set enrichment analysis) was run with GSEA version 3.0. We used msigdb ^20^ 7.3 human gene set files including: c2.cp.kegg.v7.3.symbols.gmt, c2.cp.reactome.v7.3.symbols.gmt, c5.go.bp.v7.3.symbols.gmt, h.all.v7.3.symbols.gmt as reference pathways. Produced reports were filtered for an FDR cutoff of 0.25 and used to create heatmaps.

#### Mass Spectrometry

total protein was extracted from tissue biopsies and proteomic analysis conducted by the Mass Spectrometry Proteomics Core, Advanced Technology Core at Baylor College of Medicine. Analysis was conducted on a commercial software platform, Advaita Bioinformatics and used to generate volcano plots, heatmaps, gene ontology analysis and pathway analysis.

#### Tissue Proteome Profiling

The frozen patients’ tissue was pulverized by mechanical force and 10 mg of powered samples was lysis in 50mM Ammonium bicarbonate, 1 mM CaCl_2_ by 3 min of sonication. The protein concentration of lysate was measured by Bradford method and 25 ug of lysate were digested using 0.5 ug of trypsin for 12 hours at 37 °C. Tryptic peptides were separated in a reverse-phase C18 column in a pipet tip. Peptides were eluted and separated into fifteen fractions using a stepwise gradient of increasing acetonitrile (2, 4, 6, 8, 10, 12, 14, 16, 18, 20, 22, 24, 26, 28, 30% Acetonitrile) at pH 10 then combined to five fractions (2+12+12, 4+14+24, 6+16+26, 8+18+28, 10+20+30) and vacuum dried. The dried peptide samples were resuspended with 5% methanol, 0.1% formic acid in water and analyzed on Orbitrap Fusion mass spectrometers (Thermo Fisher Scientific) coupled with an Easy-nLC 1000 nanoflow LC system (Thermo Fisher Scientific) using an in-housed trap column packed with 1.9 μm Reprosil-Pur Basic C18 beads (2 cm × 100 μm) and a 5 cm × 150 μm capillary separation column packed with 1.9 μm Reprosil-Pur Basic C18 beads with 75-min discontinuous gradient of 4–26% acetonitrile, 0.1% formic acid at a flow rate of 800 nl/min. The instrument was operated under the control of Xcalibur software version 4.1 (Thermo Fisher Scientific) in data-dependent mode, acquiring fragmentation spectra of the top 30 strongest ions. Parent mass spectrum was acquired in the Orbitrap with a full MS range of 300–1400 m/z at the resolution of 120,000. Higher-energy collisional dissociation (HCD) fragmented MS/MS spectrum was acquired in ion-trap with rapid scan mode. The MS/MS spectra were searched against the target-decoy Human RefSeq database (release Jan. 21, 2020, containing 80,872 entries) in Proteome Discoverer 2.1 interface (Thermo Fisher) with Mascot algorithm (Mascot 2.4, Matrix Science). The precursor mass tolerance of 20 ppm and fragment mass tolerance of 0.5 Da was allowed. Two maximum missed cleavage, and dynamic modifications of acetylation of N-term and oxidation of methionine were allowed. Assigned peptides were filtered with a 1% false discovery rate (FDR) using Percolator validation based on q-value. The Peptide Spectrum Matches (PSMs) output from PD2.1 was used to group peptides onto gene level using ‘gpGrouper’ algorithm ^21^. An in-house program, gpGrouper, uses a universal peptide grouping logic to accurately allocate and provide MS1 based quantification across multiple gene products. Gene-protein products (GPs) quantification was performed using the label-free, intensity-based absolute quantification (iBAQ) approach and then normalized to FOT (a fraction of the total protein iBAQ amount per experiment). FOT was defined as an individual protein’s iBAQ divided by the total iBAQ of all identified proteins within one experiment.

## RESULTS

### Echocardiography Analysis

We analyzed a total of nine parameters based on 2D Doppler Echocardiography (**Figure 1**) and only two parameters were statistically significant, LVOT peak gradient (**Figure 1E**) and LVOT mean gradient (**Figure 1F**). The LVOT peak gradient was found to be 51.20 +/- 11.9 mmHg (n=8) for non-aggressive DSS phenotype vs 76.71 +/- 13.19 mmHg (n=8) for aggressive DSS phenotype. The LVOT mean gradient was found to be 25.75 +/- 7.27 mmHg (n=8) for non-aggressive DSS phenotype vs 40.57 +/- 6.75 mmHg (n=8).

**Figure 1.**
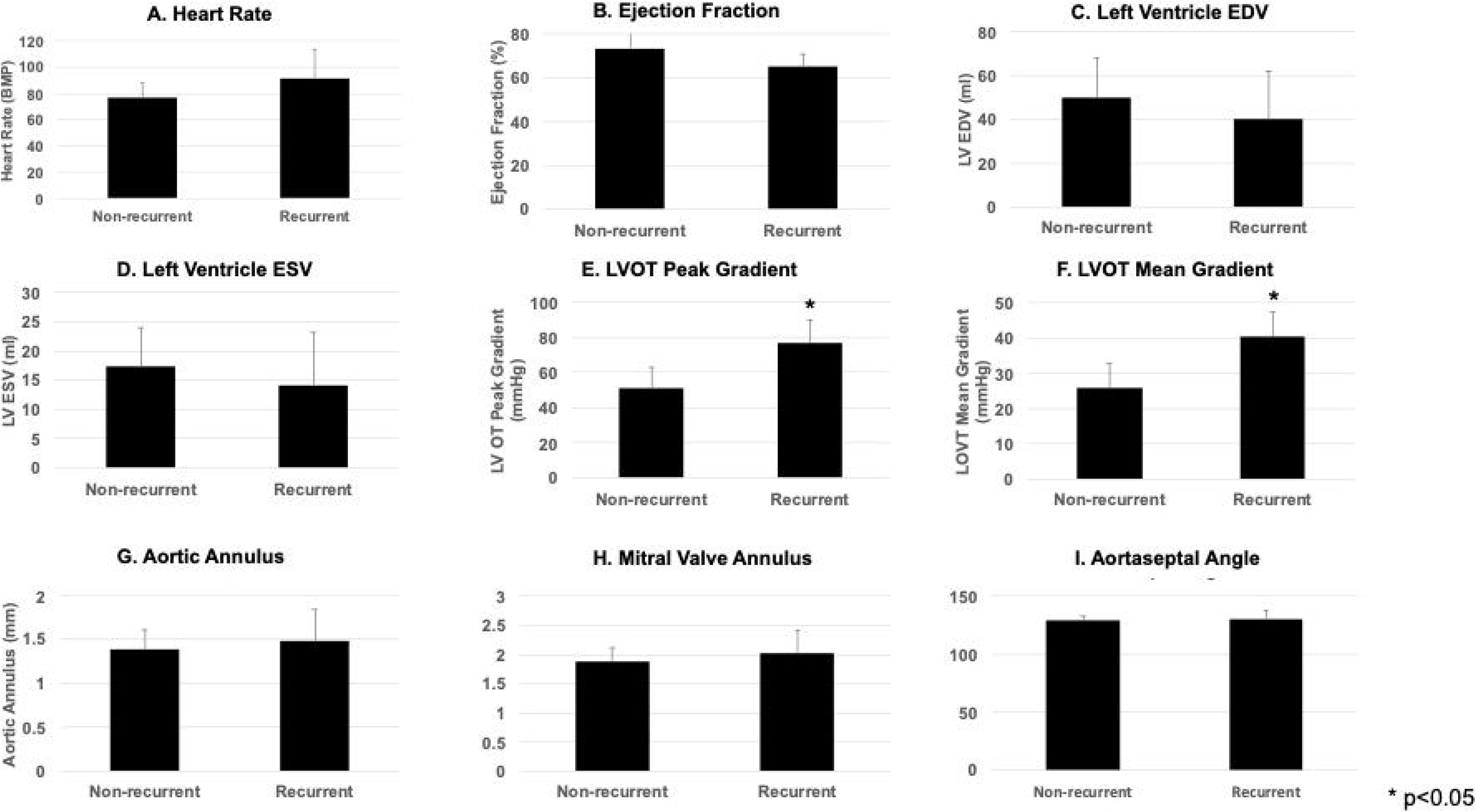
Echocardiographic Analysis of Aggressive vs Non-Aggressive DSS Patients. Patient data was analyzed for the following parameters, with N=8 per group: (A) Heart Rate, (B) Ejection Fraction, (C) LVEDV, (D) LVESV, (E) LVOT Peak Gradient, (F) LVOT Mean Gradient, (G) Aortic Annulus, (H) Mitral Valve Annulus, (I) Aortaseptal Angle. N=8 per group, *p<0.05.

### Genomic Analysis for AS1 vs NA Surgery Based on Volcano Plots

the volcano plot (**Figure 2A**) shows the differential expression of genes in AS1 vs NA samples, with 11,794 genes downregulated and 1,388 genes upregulated and no change in 1,821 genes. The full list of genes, along with the fold changes is presented in **Table 1, supplemental data,** and the top 100 genes based on significance is presented in **Table 2, supplemental data,** and genes relevant to DSS are presented in **Table 1**. Many of the significant genes listed in Table are known to be associated with extracellular matrix (ECM) function and cellular interactions with the ECM. While **Table 1** list 25 genes relevant to DSS, 15 genes are known to regulate ECM function and subsequent cellular function. Modulation of actin, elastin and hyaluronic acid are shown to be important modulators of DSS through changes in gelsolin, ficolin 1 and hyaluronan synthase 2 genes.

**Figure 2.**
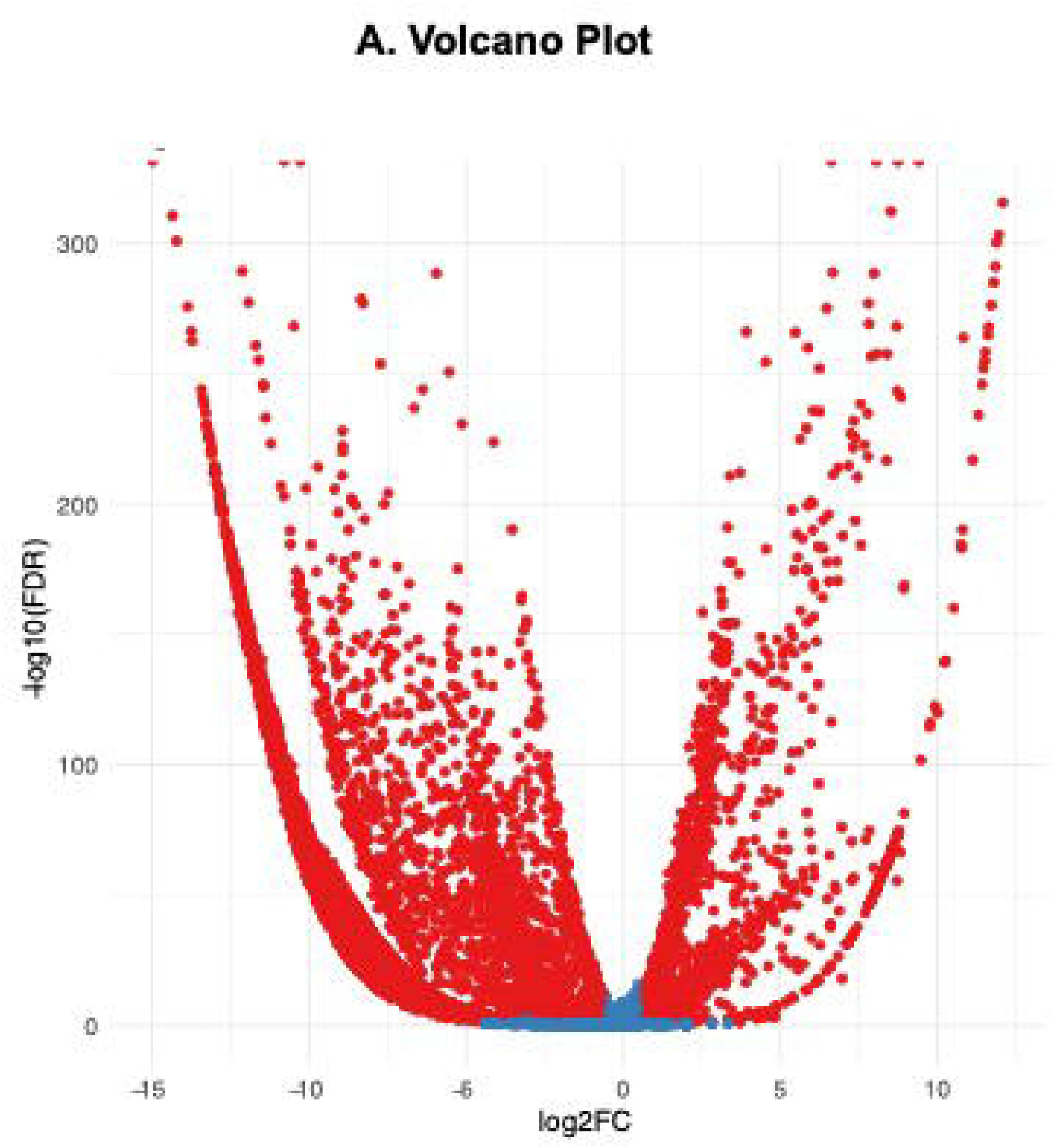
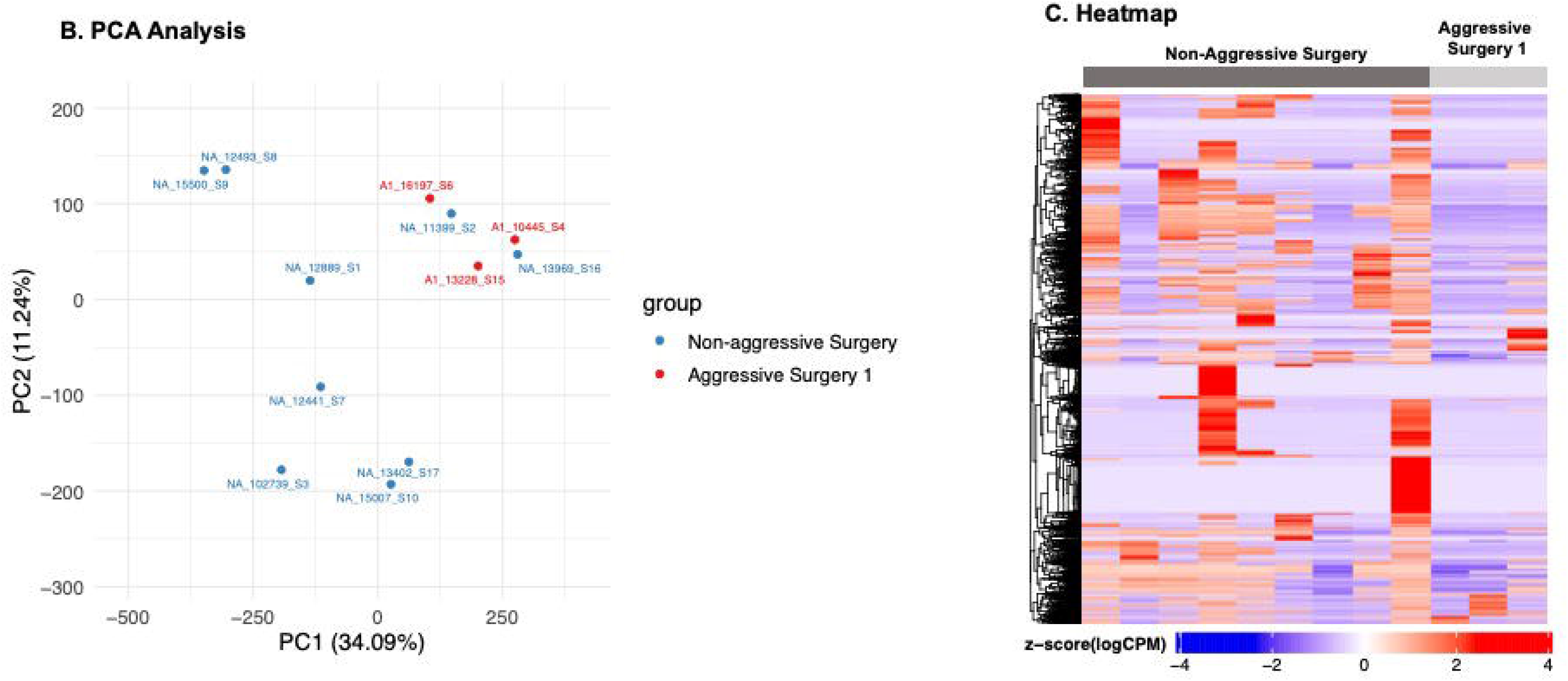
Differential Gene Expression in Aggressive vs Non-Aggressive DSS Patients. **(A) Volcano Plot** – 11794 genes were downregulated, and 1388 genes upregulated, non-aggressive DSS patients relative to aggressive DSS patients. **(B) PCA Analysis** – distinct separation of aggressive (A) vs non-aggressive patients. **(C) Heatmap** – shows significant changes in gene expression of aggressive DSS patients relative to non-aggressive DSS patients.

**Table 1.**
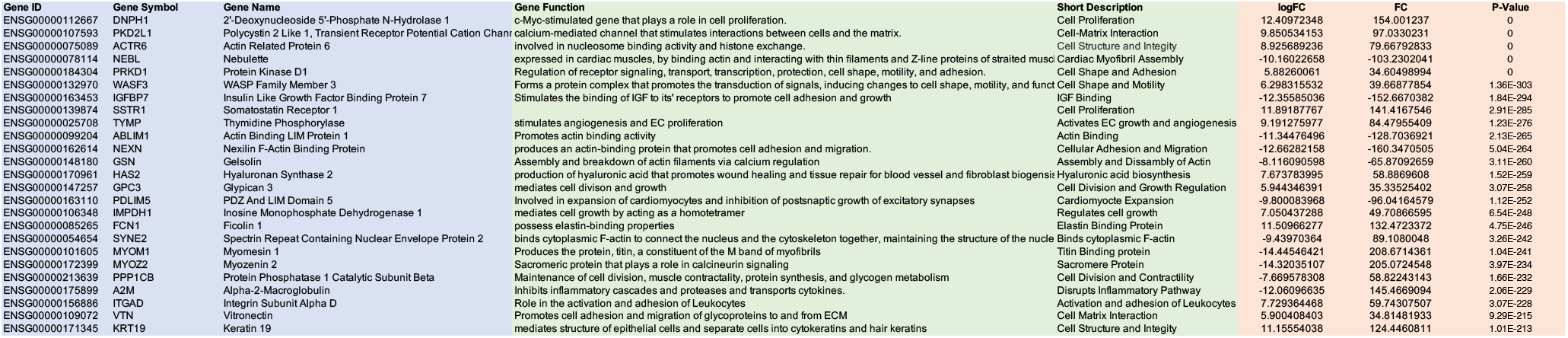
List of Differentially Regulated Genes for AS1 vs NA Relevant for DSS.

**Table 2.**
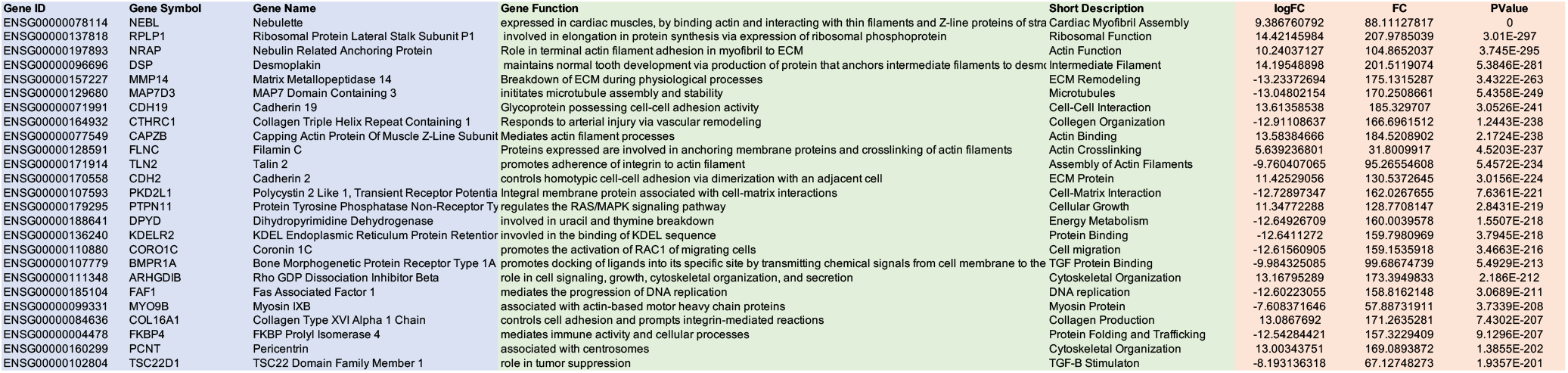
List of Differentially Regulated Genes for AS2 vs AS1 Relevant for DSS.

### PCA Analysis and Heatmaps for AS1 vs NA Surgery

there was clear separation of the two groups when placed on PCA plot, with AS1 samples grouping on the top right of the plot while samples from NA surgery grouping to the lower portions of the PCA plot (**Figure 2B**). Plotting differential gene expression in the form of a heatmaps showed significant changes in gene expression between AS1 and NA samples (**Figure 2C**).

### Genomic Analysis of DSS Patients with Aggressive Phenotype Based on Volcano Plots

the volcano plot for DSS patients with an aggressive phenotype (**Figure 3A**) demonstrated a decrease in 11,422 genes and an increase in 1,666 genes in patients at the time of AS1 vs AS2. The full list of genes, along with the fold changes is presented in **Table 3, supplemental data,** and the top 100 genes based on significance is presented in **Table 4, supplemental data,** and genes relevant to DSS are presented in **Table 2**. Many of the genes that show significant changes between AS2 relative to AS1 include genes involved in ECM function and cellular interaction with the ECM. Specific examples from **Table 2** include genes involved in actin remodeling (NRAP, CAPZB, FLNC, TLN2), collagen organization (CTHRC1, COL16A1), ECM structure and organization (DSP, ARHGDIB, PCNT) and cell-matrix interaction (PKD2L1).

**Figure 3.**
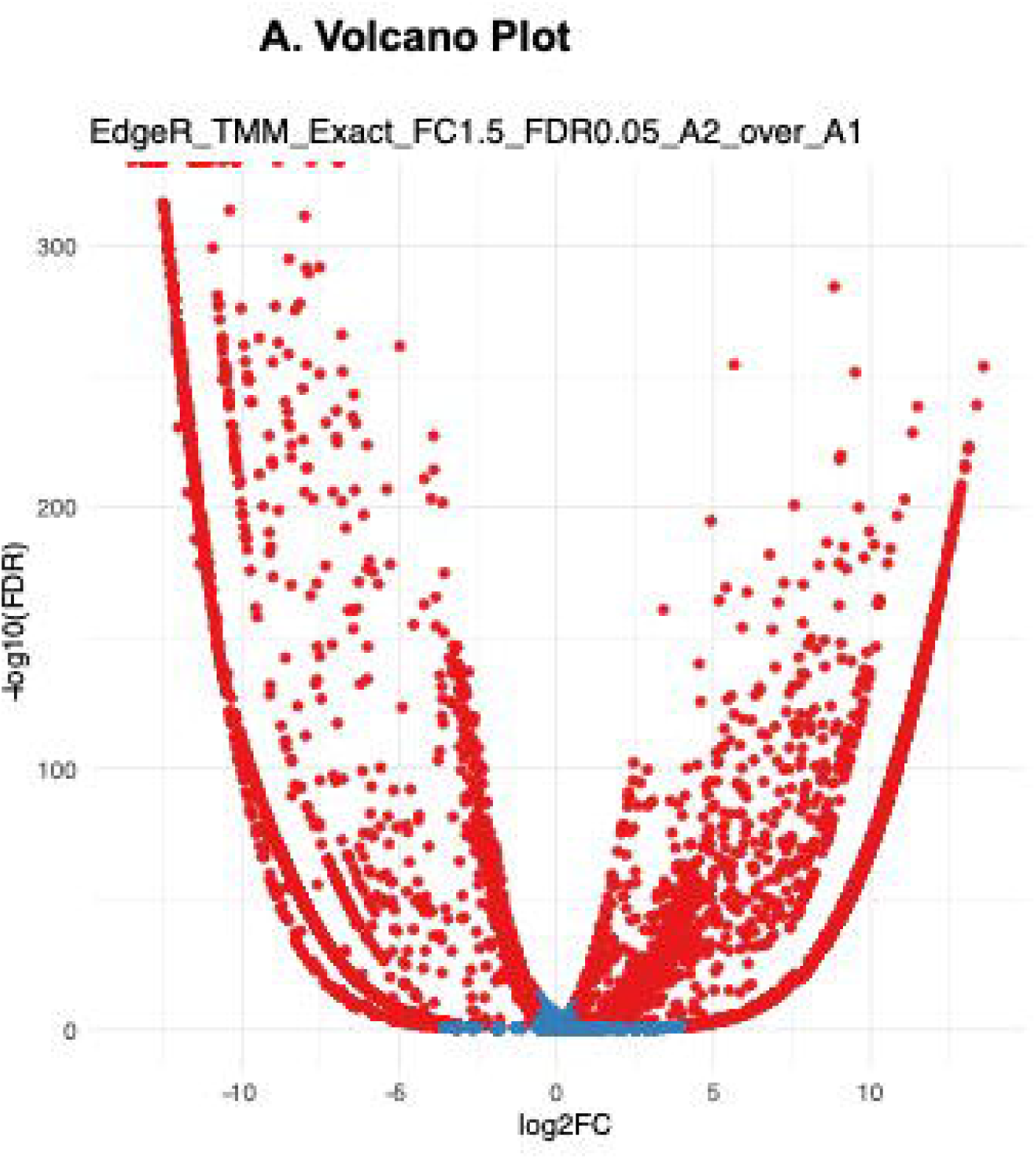
Differential Gene Expression in DSS Patients at the time of Aggressive Surgery 1 vs Aggressive Surgery 2. **(A) Volcano Plot** – 11,422 genes were downregulated, and 1,666 genes upregulated in patients at the time of first surgery relative to second surgery for DSS patients with aggressive DSS phenotype. **(B) PCA Analysis** – clustering of samples from aggressive DSS patients at the time of first and second surgery. There was no separation of these two groups into distinct populations on the PCA plot. **(C) Heatmap** – changes in the gene expression between AS1 vs AS2 were minimal, as illustrated on the heatmap based on the intensity of the color changes.

**Table 3.** List of Differentially Regulated Peptides for AS1 vs NA Relevant for DSS.

| Gene Symbol | Gene Function | Short Description | Entrez | Log FC | Fold Change | Adjusted P-Value |
| --- | --- | --- | --- | --- | --- | --- |
| HAPLN4 | ECM formation | ECM formation | 404037 | -10 | 100 | 3.04545E-06 |
| TMSB4X | regulates actin polymerization | actin polymerization | 7114 | 10 | 100 | 1.02451E-05 |
| UBE2K | regulates degradation of short-lived proteins | degradation of abnormal proteins | 3093 | -7.7338444 | 59.81234967 | 1.03846E-05 |
| KRT6C | role in the structure of epithelial cells | structural integrity of epithelial cells | 286887 | -10 | 100 | 1.12413E-05 |
| TMSB10 | cytoskeleton organization | cytoskeleton organization | 9168 | 10 | 100 | 2.79613E-05 |
| NCKAP1 | actin filament remodeling | actin filament reorganization | 10787 | 8.02164529 | 64.34679323 | 4.27585E-05 |
| CHMP1A | role in sorting proteins of lysosomes required for transport complex III | protein sorting | 5119 | 7.9560239 | 63.29831634 | 0.000223835 |
| FETUB | regulates insulin and hepatocyte growth factor receptors and responds to systemic inflammation | protease inhibitor | 26998 | -8.9542525 | 80.1786379 | 0.000228621 |
| CNN2 | structural integrity of actin filaments | actin filament organization | 1265 | -10 | 100 | 0.000279096 |
| ITGB5 | cell adhesion and cell-surface signaling | cell adhesion and cell-surface sigr | 3693 | 8.45477738 | 71.48326052 | 0.000304806 |
| KRT8 | heteropolymerization of type I and II keratins to form intermediate filaments in epithelial cytoplasm | intermediate filament construction | 3856 | -10 | 100 | 0.001052125 |
| TNXB | organization and maintenance of tissues | organizes and maintains various ti | 7148 | -1.9755827 | 3.902927109 | 0.010365286 |
| ACTA2 | production of actin proteins | actin protein production | 59 | 1.11404773 | 1.241102353 | 0.01939829 |
| SH3BGRL | scaffold protein | scaffold protein | 6451 | -3.1864835 | 10.15367691 | 0.023649103 |
| ACTN1 | actin-binding activity | actin binding activity | 87 | 1.83744669 | 3.376210353 | 0.024906938 |
| KRT7 | actin binding activity | actin binding activity | 3855 | -7.1514309 | 51.14296458 | 0.025211992 |
| MAP2 | microtubule assembly | microtubule assembly | 4133 | -3.842916 | 14.76800342 | 0.036005958 |
| TBCB | regulates the dissociation of tubulin heterodimers | dissociation of tubulin heterodimer | 1155 | -6.1917647 | 38.33795013 | 0.036796979 |
| COL3A1 | production of type III collagen | production of collagen | 1281 | -1.4281377 | 2.039577308 | 0.037296285 |
| WASF2 | links receptor kinases and actin | multi-protein complex | 10163 | -4.2202253 | 17.81030121 | 0.038276362 |
| LRG1 | protein-protein interaction, signal transduction, cell-adhesion, and cell development | protein-protein interaction | 116844 | -5.0961303 | 25.97054403 | 0.039337959 |
| CDC42EP4 | regulates actin cytoskeleton organization | regulates actin cytoskeleton orgar | 23580 | -3.5684455 | 12.73380329 | 0.045674892 |

**Table 4.**
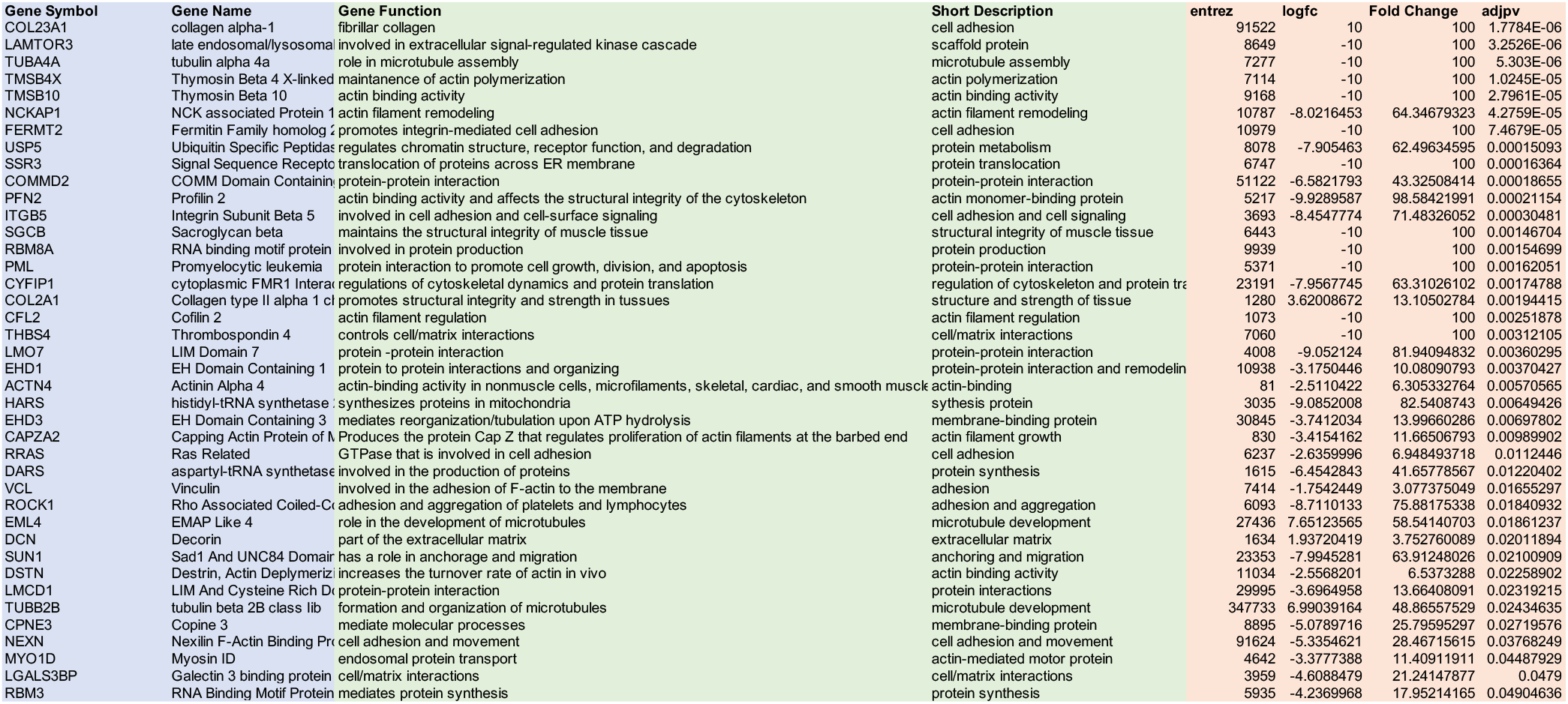
List of Differentially Regulated Peptides for AS2 vs AS1 Relevant for DSS.

### PCA Analysis and Heatmaps for DSS Patients with Aggressive Phenotype

the PCA plot (**Figure 3B**) shows close clustering of samples from first and second DSS surgery, with very little separation of the samples. This is further illustrated in the heatmap (**Figure 3C**), which shows very little changes in the differential expression of genes, observed by the flat color scheme, at the time of first DSS surgery, relative to the second surgery.

### Pathway Analysis Based on Genomic Analysis

pathway analysis based on Hallmark pathways (**Figure 4A**) and **Table 5**, **supplemental data**, show significant upregulation of epithelial to mesenchymal transition. Additional significant pathways based on Hallmark pathways show upregulation in pathways related to ECM function and remodeling, to include TGF-beta signaling, apical junction and protein secretion (**Figure 4A**). Similar results were observed by analysis of the KEGG pathways (**Figure 4B**) and **Table 6**, **supplemental data**, with significant increase in pathways promoting ECM function, remodeling and cell-matrix interactions in DSS patients with an aggressive phenotype when compared with those with a non-aggressive phenotype, both at the time of first surgery. Specific examples of KEGG pathways that were upregulated in aggressive DSS patients included adherens junction, focal adhesion, protein export, tight junction, TGF-beta signaling, VEGF signaling and MAPK signaling pathways (**Figure 4B**).

**Figure 4.**
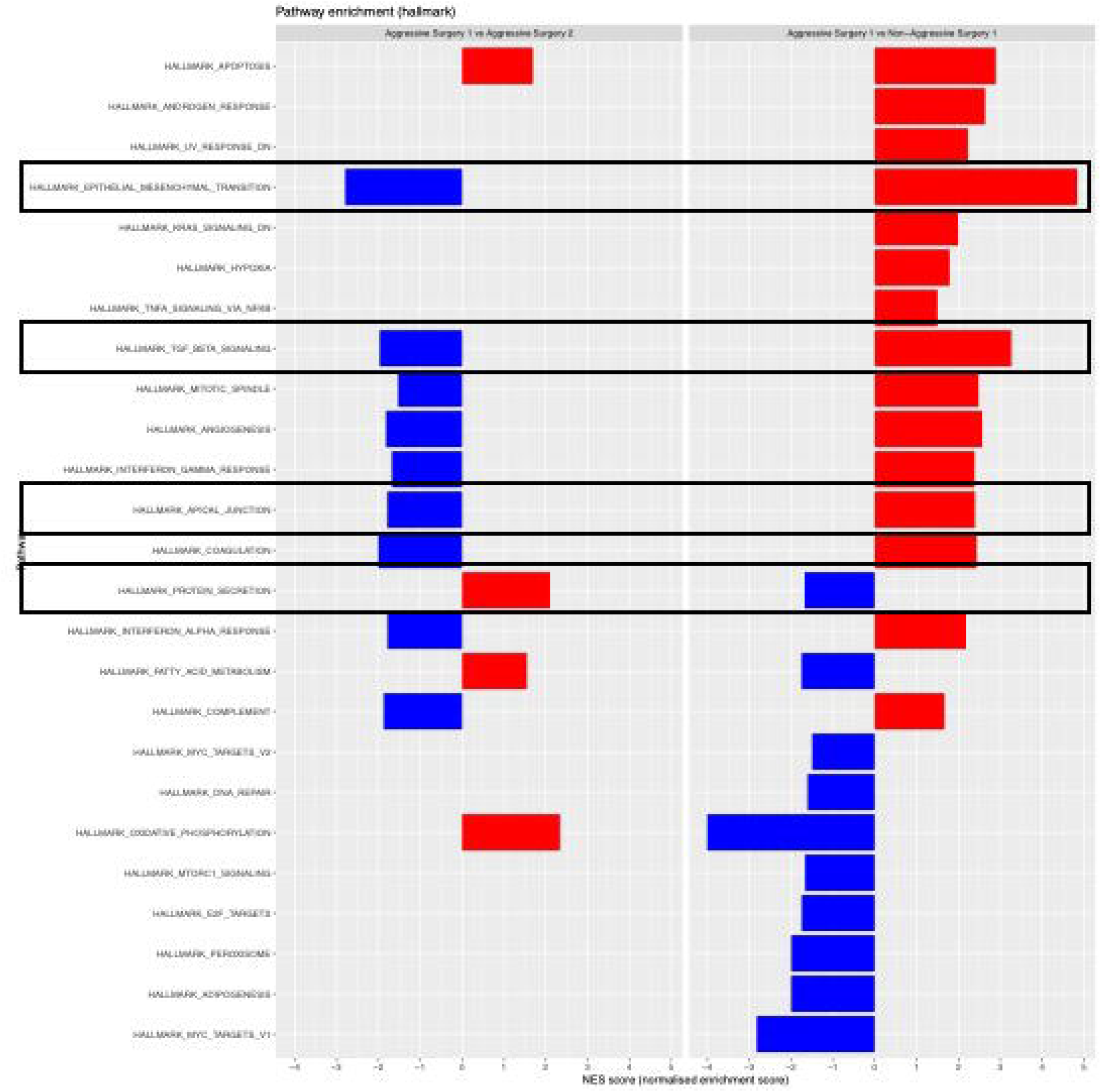
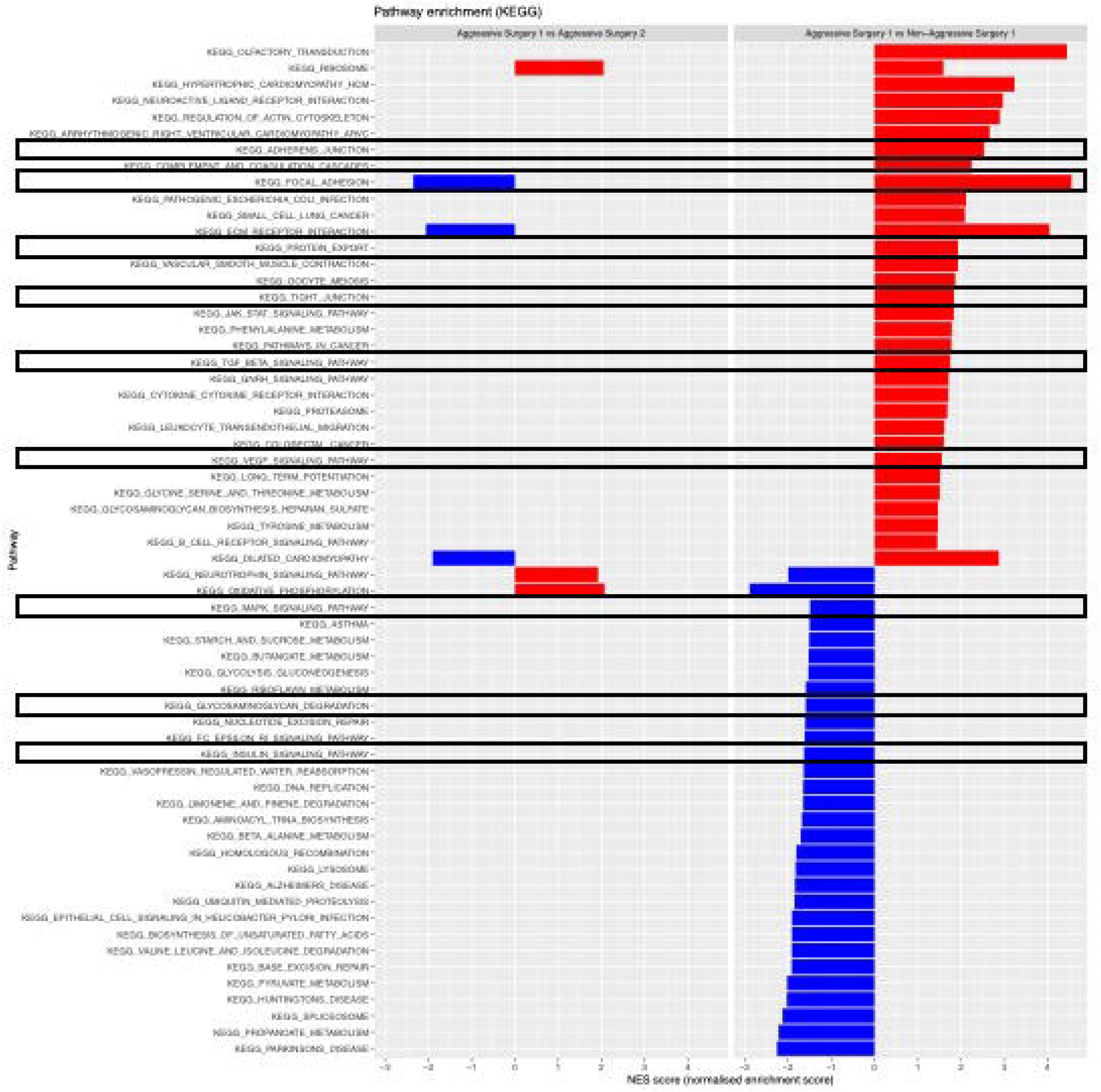
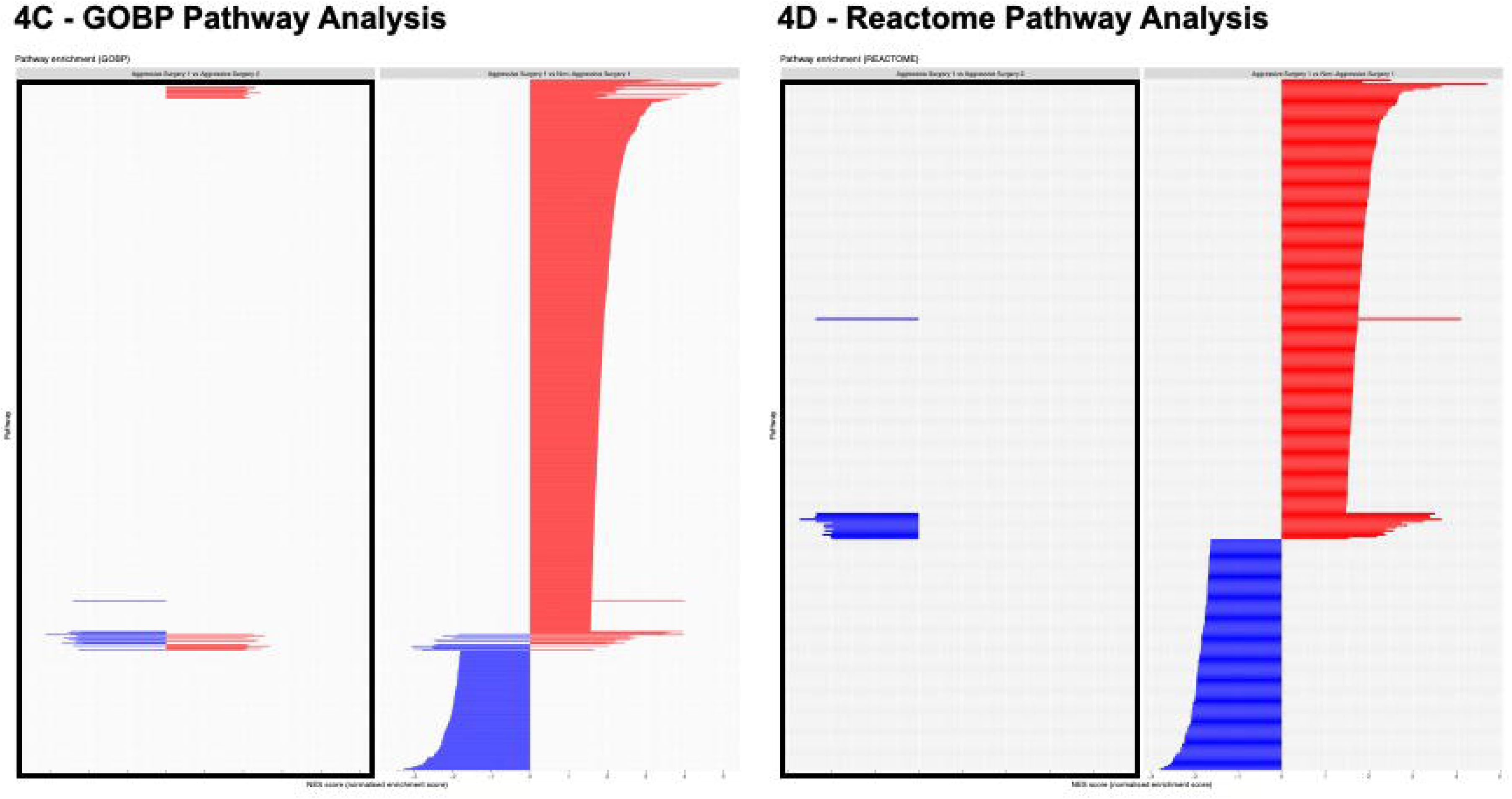
Hallmark, KEGG, GOPB and Reactome Pathways. Differential expression of **(A) Hallmark, (B) KEGG, (C) GOBP and (D) Reactome Pathways** in DSS Patients with aggressive phenotype comparing samples at first and second surgery (left panel) and samples at the time of first surgery, comparing aggressive vs non-aggressive (right panel). For the **Hallmark (A)** and **KEGG (B)** Pathways, the boxes show differential expression of pathways relevant to DSS. For **GOBP (C)** and **Reactome (D)** Pathway, the box shows very little changes in differential expression of all pathways in DSS patients with aggressive phenotype, when comparing second surgery relative to first surgery.

**Table 5.** List of Differentially Regulated Pathways Based on Gene Ontology Analysis for AS1 vs NA Relevant for DSS.

| Gene Ontology ID | Gene Ontology Name | Differential Enriched Proteins | All Proteins | P-Value |
| --- | --- | --- | --- | --- |
| GO:0060249 | anatomical structure homeostasis | 11 | 127 | 0.00116 |
| GO:0030036 | actin cytoskeleton organization | 17 | 303 | 0.00705 |
| GO:0007010 | cytoskeleton organization | 24 | 491 | 0.0078 |
| GO:0042989 | sequestering of actin monomers | 2 | 5 | 0.00822 |
| GO:0030029 | actin filament-based process | 18 | 345 | 0.0116 |
| GO:0030810 | positive regulation of nucleotide biosynthetic process | 2 | 6 | 0.0121 |
| GO:0007015 | actin filament organization | 12 | 199 | 0.01372 |
| GO:0030199 | collagen fibril organization | 4 | 34 | 0.01712 |
| GO:0051127 | positive regulation of actin nucleation | 2 | 8 | 0.02172 |
| GO:0051258 | protein polymerization | 9 | 140 | 0.02184 |
| GO:0090066 | regulation of anatomical structure size | 10 | 170 | 0.02781 |
| GO:0060940 | epithelial to mesenchymal transition involved in cardiac fibroblast development | 1 | 1 | 0.02967 |
| GO:1901202 | negative regulation of extracellular matrix assembly | 1 | 1 | 0.02967 |
| GO:1904527 | negative regulation of microtubule binding | 1 | 1 | 0.02967 |
| GO:2000575 | negative regulation of microtubule motor activity | 1 | 1 | 0.02967 |
| GO:0051493 | regulation of cytoskeleton organization | 12 | 223 | 0.03078 |
| GO:0060491 | regulation of cell projection assembly | 5 | 60 | 0.03136 |
| GO:1900026 | positive regulation of substrate adhesion-dependent cell spreading | 3 | 24 | 0.03267 |
| GO:0008360 | regulation of cell shape | 5 | 61 | 0.03338 |
| GO:0032970 | regulation of actin filament-based process | 10 | 177 | 0.03543 |
| GO:0048771 | tissue remodeling | 4 | 44 | 0.04003 |
| GO:0031589 | cell-substrate adhesion | 9 | 156 | 0.04022 |
| GO:0032956 | regulation of actin cytoskeleton organization | 9 | 156 | 0.04022 |
| GO:0032271 | regulation of protein polymerization | 7 | 110 | 0.04328 |
| GO:0032879 | regulation of localization | 30 | 751 | 0.04369 |
| GO:0009605 | response to external stimulus | 29 | 723 | 0.04531 |
| GO:0030038 | contractile actin filament bundle assembly | 4 | 46 | 0.04602 |
| GO:0043149 | stress fiber assembly | 4 | 46 | 0.04602 |
| GO:0034105 | positive regulation of tissue remodeling | 2 | 12 | 0.04741 |
| GO:0034315 | regulation of Arp2/3 complex-mediated actin nucleation | 2 | 12 | 0.04741 |
| GO:0030041 | actin filament polymerization | 6 | 90 | 0.04933 |
| GO:0051494 | negative regulation of cytoskeleton organization | 5 | 68 | 0.04977 |

**Table 6.** List of Differentially Regulated Pathways Based on Gene Ontology Analysis for AS2 vs AS1 Relevant for DSS.

| Gene Ontology ID | Gene Ontology Name | Differentially Enriched Proteins | All Proteins | P-Value |
| --- | --- | --- | --- | --- |
| GO:0007015 | actin filament organization | 18 | 189 | 0.0001 |
| GO:0007010 | cytoskeleton organization | 32 | 469 | 0.00016 |
| GO:0090066 | regulation of anatomical structure size | 16 | 161 | 0.00016 |
| GO:0030036 | actin cytoskeleton organization | 23 | 291 | 0.0002 |
| GO:0032956 | regulation of actin cytoskeleton organization | 15 | 149 | 0.00022 |
| GO:0099515 | actin filament-based transport | 4 | 10 | 0.00028 |
| GO:0032970 | regulation of actin filament-based process | 16 | 169 | 0.00028 |
| GO:0030029 | actin filament-based process | 24 | 332 | 0.00055 |
| GO:0008154 | actin polymerization or depolymerization | 11 | 101 | 0.00084 |
| GO:0097435 | supramolecular fiber organization | 23 | 323 | 0.0009 |
| GO:0030832 | regulation of actin filament length | 10 | 87 | 0.00095 |
| GO:0030031 | cell projection assembly | 13 | 146 | 0.00192 |
| GO:0034315 | regulation of Arp2/3 complex-mediated actin nucleation | 3 | 9 | 0.00324 |
| GO:0110053 | regulation of actin filament organization | 11 | 120 | 0.00342 |
| GO:0007043 | cell-cell junction assembly | 6 | 44 | 0.00438 |
| GO:0060249 | anatomical structure homeostasis | 11 | 124 | 0.00441 |
| GO:0048771 | tissue remodeling | 6 | 45 | 0.0049 |
| GO:0030833 | regulation of actin filament polymerization | 8 | 75 | 0.00494 |
| GO:0030043 | actin filament fragmentation | 2 | 4 | 0.00732 |
| GO:2000601 | positive regulation of Arp2/3 complex-mediated actin nucleation | 2 | 4 | 0.00732 |
| GO:0098609 | cell-cell adhesion | 15 | 210 | 0.00739 |
| GO:0045185 | maintenance of protein location | 5 | 35 | 0.00753 |
| GO:0030041 | actin filament polymerization | 8 | 83 | 0.00909 |
| GO:0051125 | regulation of actin nucleation | 3 | 13 | 0.00993 |
| GO:1902903 | regulation of supramolecular fiber organization | 12 | 159 | 0.01076 |
| GO:0030836 | positive regulation of actin filament depolymerization | 2 | 6 | 0.01745 |
| GO:0051127 | positive regulation of actin nucleation | 2 | 6 | 0.01745 |
| GO:0030048 | actin filament-based movement | 7 | 76 | 0.01819 |
| GO:0034330 | cell junction organization | 14 | 213 | 0.01902 |
| GO:1902905 | positive regulation of supramolecular fiber organization | 8 | 96 | 0.02074 |
| GO:0030838 | positive regulation of actin filament polymerization | 5 | 45 | 0.02132 |
| GO:1901881 | positive regulation of protein depolymerization | 2 | 7 | 0.02387 |
| GO:2000810 | regulation of bicellular tight junction assembly | 2 | 7 | 0.02387 |
| GO:0051495 | positive regulation of cytoskeleton organization | 8 | 99 | 0.02449 |
| GO:0070830 | bicellular tight junction assembly | 3 | 18 | 0.02486 |
| GO:0030837 | negative regulation of actin filament polymerization | 4 | 32 | 0.02617 |
| GO:0032271 | regulation of protein polymerization | 8 | 102 | 0.0287 |
| GO:0120192 | tight junction assembly | 3 | 19 | 0.02877 |
| GO:0120193 | tight junction organization | 3 | 19 | 0.02877 |
| GO:0045216 | cell-cell junction organization | 6 | 66 | 0.02972 |
| GO:0150105 | protein localization to cell-cell junction | 2 | 8 | 0.03108 |
| GO:0150106 | regulation of protein localization to cell-cell junction | 1 | 1 | 0.03591 |
| GO:1904702 | regulation of protein localization to adherens junction | 1 | 1 | 0.03591 |
| GO:0051014 | actin filament severing | 2 | 9 | 0.03904 |
| GO:0034329 | cell junction assembly | 9 | 130 | 0.04217 |
| GO:0001954 | positive regulation of cell-matrix adhesion | 3 | 22 | 0.0423 |

Analysis of GOBP and Reactome pathways showed very small changes in the differential expression of all pathways in DSS patients with an aggressive phenotype, at the time of second surgery compared with the first surgery (**Figure 4C and 4D**) and **Table 7** and **Table 8**, **supplemental data**.

**Table 7.** List of Differentially Regulated Pathways Based on Pathway Analysis for AS1 vs NA Relevant for DSS.

| Pathway Name | Differentially Enriched Proteins | Total Proteins | P-Value | pAcc | pComb | pORA |
| --- | --- | --- | --- | --- | --- | --- |
| Regulation of actin cytoskeleton | 10 | 111 | 0.00912024 | 0.87456272 | 0.00912024 | 0.0013466 |
| Adherens junction | 5 | 33 | 0.00942502 |  |  | 0.00244601 |
| Focal adhesion | 9 | 110 | 0.01827056 | 0.5812094 | 0.01827056 | 0.00452978 |

**Table 8.**
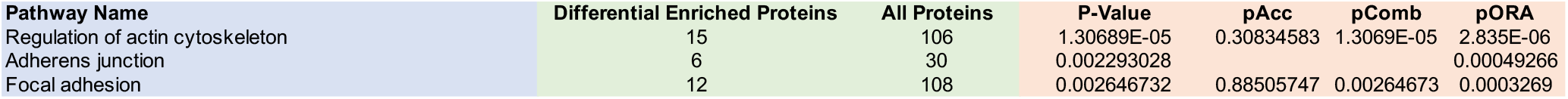
List of Differentially Regulated Pathways Based on Pathway Analysis for AS2 vs AS1 Relevant for DSS.

| Pathway Name | Differential Enriched Proteins | All Proteins | P-Value | pAcc | pComb | pORA |
| --- | --- | --- | --- | --- | --- | --- |
| Regulation of actin cytoskeleton | 15 | 106 | 1.30689E-05 | 0.30834583 | 1.3069E-05 | 2.835E-06 |
| Adherens junction | 6 | 30 | 0.002293028 |  |  | 0.00049266 |
| Focal adhesion | 12 | 108 | 0.002646732 | 0.88505747 | 0.00264673 | 0.0003269 |

### Differential Expression of Peptides for DSS Patients Based on Volcano Plots

In the case of AS1 vs NA, a total of 3714 peptides were detected, of which 110 were significant and 63 were downregulated and 47 were upregulated, **Figure 5A** and **Table 9**, **supplemental data**. A list of the top 100 peptides based on significance are listed in **Table 10**, **supplemental data** and the peptides most relevant to DSS are listed in **Table 3**. A total of 22 peptides were identified as significant for DSS, all of which were involved in ECM assembly, function and interaction with cells, **Table 3**. Of the 22 peptides relevant to DSS, 7 (or 31.8%) were involved in actin function and assembly (TMSB4X, NCKAP1, CNN2, ACTA2, ACTN1, KRT7, CDC42EP4).

**Figure 5.**
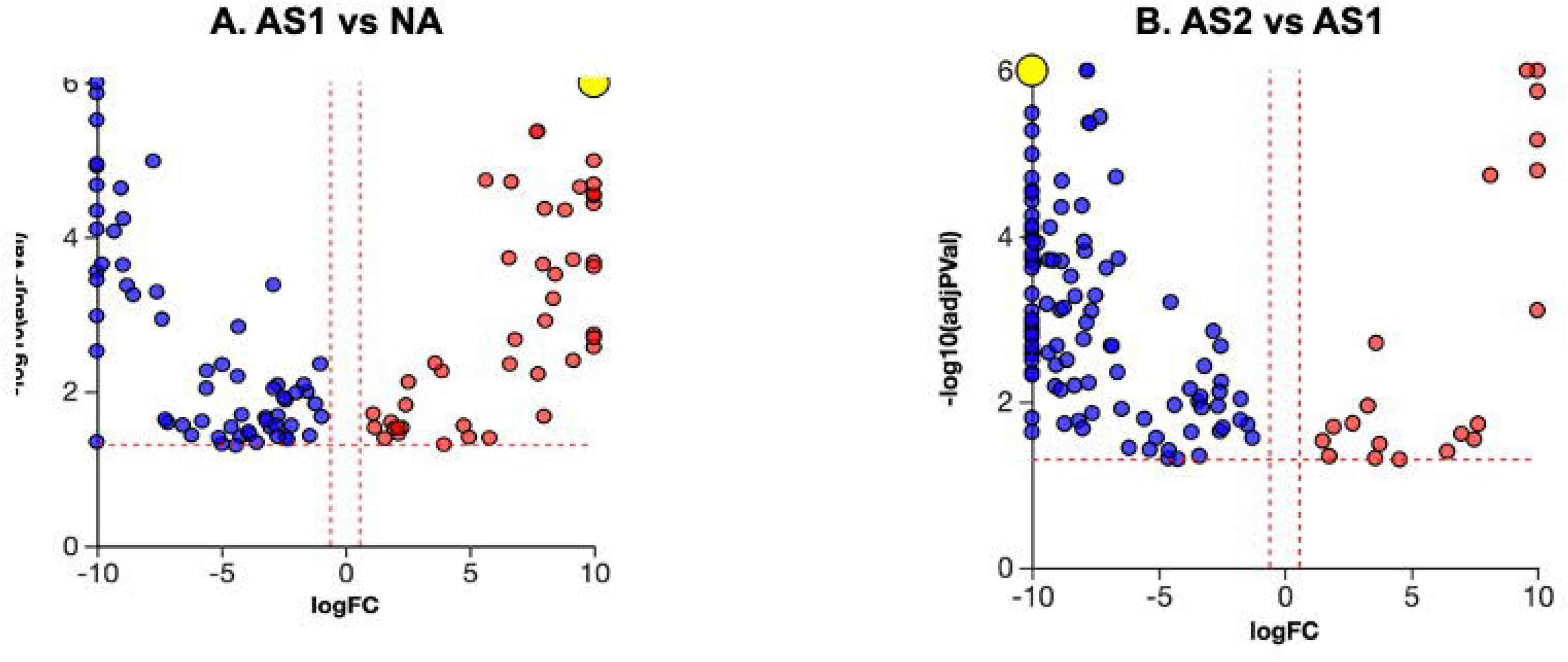
Volcano Plots Showing Differential Protein Expression Based on Mass Spectrometry. **(A) AS1 vs NA** – 110 peptides were noted to be significant (p<0.05), with 63 downregulated (blue) and 47 upregulated peptides (red). **(B) AS2 vs AS1** – 126 peptides were noted to be significant (p<0.05), with 106 downregulated (blue) and 20 upregulated peptides (red).

In the case of AS2 vs AS1, a total of 3590 peptides were differentially regulated, of which 126 were significant (p<0.05), 106 peptides downregulated, and 20 peptides upregulated, **Figure 5B**, **Table 11**, **supplemental data**. A list of the top 100 peptides based on significance are listed in **Table 12**, **supplemental data** and the peptides most relevant to DSS are listed in **Table 4**. As can be seen from **Table 4**, a total of 40 peptides were determined to be relevant for DSS and all the peptides were involved in ECM assembly, function, remodeling and attachment and anchoring of cells to the ECM. Of the 40 peptides found relevant for DSS, 9 or 22.5% were involved in process related to actin assembly and function (TMSB4X, TMSB10, NCKAP1, PFN2, CFL2, ACTN4, CAPZA2, DSTN, MYO1D).

### Gene Ontology Analysis for Differentially Enriched Peptides Based on Mass Spectrometry Data

**Table 13**, **supplemental data**, lists all the pathways that were differentially regulated in DSS patients at the of first surgery, when an aggressive phenotype was compared to a non-aggressive phenotype. As shown in **Table 13**, **supplemental data**, a total of 2856 pathways were differentially regulated, although only 405 were statistically significant (p<0.05) and only 32 relevant for DSS, **Table 5**. The key pathways identified by gene ontology analysis were based on epithelial to mesenchymal transition and 31 pathways known to regulate ECM assembly and function with 10 of 32 (or 31.25%) of the pathways focused on actin assembly and function. Of the 10 pathways focused on actin remodeling, 6 pathways are presented in **Figure 5** and include critical individual peptide components that are known to regulate these pathways. For comparison, the changes in these pathways for DSS patients with an aggressive phenotype, at the time of second surgery compared with first surgery, is also presented in **Figure 5**. This comparison serves to illustrate the key point that pathways that are differentially regulated at the time of first surgery, continue to be further downregulated at the time of second surgery, when compared against first surgery in DSS patients with aggressive DSS phenotype. **Table 14**, **supplemental data**, lists all the pathways that are differentially regulated based on gene ontology analysis for AS2 vs AS1 and **Table 6**, list the pathways that are relevant for DSS. As noted in **Table 6**, a total of 46 pathways were differentially regulated based on gene ontology analysis, all of which were focused on ECM assembly, remodeling and cell-matrix interaction. In addition, 21 (or 45.6%) of the 46 differentially regulated pathways were related to actin assembly and function.

### Regulation of Actin Cytoskeleton Based on Proteomics Data

A total of 160 pathways were differentially regulated in DSS patients at the time of first surgery, for patients with an aggressive vs non-aggressive phenotype, **Table 15**, **supplemental data**. Three differentially regulated pathways were determined to be relevant to DSS, **Table 7**, and included the regulation of actin cytoskeleton, pathways related to adherens junctions and pathways related to focal adhesions, of which are known to be key regulators in the formation of ECM.

**Figure 7A** shows the points at which specific peptides were either upregulated (red) or downregulated (blue) on the pathway responsible for the production of actin cytoskeleton. As shown in **Figure 7B**, a total of 10 peptides were differentially regulated in the production of actin and these include RAC2, RAC3, TMSB4X, ITGB5, NCKAP1, PPP1CC, PPP1CB, WASF2, RAC1 and ACTN1. Furthermore, the network diagram in **Figure 7C** shows the relationship of these individual constituent peptides towards the production of actin, the net effect of which resulted in downregulation of actin in DSS patients with an aggressive phenotype, compared with non-aggressive DSS patients, both at the time of first surgery.

This was in contrast to actin production in DSS patients with an aggressive phenotype at the time of second surgery, when compared with the first surgery. As can be seen from **Table 16**, **supplemental data**, 176 pathways were differentially regulated in aggressive DSS patients at the time of surgery two relative to surgery one. As was the case for AS1 vs NA, three pathways related to ECM assembly and function were found to be relevant to aggressive DSS patients, at the time of surgery two relative to surgery one, **Table 8**: regulation of actin cytoskeleton, adherens junction and focal adhesion.

The regulation of actin cytoskeleton exhibited significant changes in aggressive DSS patients at the time of second surgery (**Figure 7D-F**). As can be seen in **Figure 7D**, there are numerous points along the pathway for the production of actin where specific peptides are downregulated (blue boxes), with no peptides being upregulated. A total of 15 peptides are upregulated in DSS patients at the time of second surgery, related to the first surgery (**Figure 7E**) and these are CFL2, PPP1CB, RAC2, RAC3, TMSB4X, PFN2, ROCK1, ITGB5, NCKAP1, CYFIP1, MYLK3, RRAS, ACTN4, MYH14, VCL. The network diagram (**Figure 7F**) shows the relationship between individual peptides and production of actin.

## DISCUSSION

Pediatric patients with DSS are faced with the uncertainty of reoccurrence of the fibrotic membrane. While reoccurrence occurs in 30% of DSS patients, all patients are managed the same post-operatively, with frequent follow-up visits to the hospital to test for formation of the fibrotic membrane for a second time. Currently, there is no method to predict the probability of reoccurrence based on genetic signatures of the pediatric patient. The ability to identify DSS patients at a high risk for reoccurrence would dramatically influence the care of these patients. High risk DSS patients would be monitored and managed more aggressively, while low risk patients can be subjected to a treatment regime with reduced frequency of follow-up care. The net result of this would be the development of treatment plans that are customized to the needs of specific DSS patient populations, categorized into those that are at high-risk vs low risk for reoccurrence of the fibrotic membrane.

In this study, we sought to test the study hypothesis “*differential expression of genes and proteins in DSS patients at the time of first surgery would predict the risk of reoccurrence after resection of the fibrotic membrane”*. In other words, we hypothesized that reoccurrence of the fibrotic membrane in DSS patients is due to differential expression at the gene and protein level. Identification of genetic markers of DSS reoccurrence is powerful and can lead to the development of a predictive lab-based test to identify the risk of reoccurrence. Based on current standard of card, the fibrotic membrane is removed and often time discarded during initial DSS surgery. Rather than discarding the fibrotic membrane, our study sought to define a method to use easily accessible tools, like RNA-sequencing and mass spectrometry, to identify the risk of reoccurrence in DSS patients. In addition, we sought to identify differential expression of genes/proteins of resected DSS membranes at the time of first and second surgery to understand the changes that drive the progression of a complex disorder like DSS.

In this study we sought to identify differentially regulated genes using RNA-sequencing and proteins, using mass spectrochemistry, for fibrotic membranes resected from DSS patients at the time of first surgery (for non-recurrent patients) and first and second surgery (for recurrent patients). Our objective was to identify specific genes, proteins and pathways that are differentially regulated between these two patient populations. The data obtained from this study will increase our understanding of the molecular regulators of DSS and the pathways that are differentially regulated in recurrent vs non-recurrent DSS patients. The potential implications of this data extend beyond as modulation of these pathways can lead to potential therapies and limit the progression of DSS.

We first sought to evaluate cardiac hemodynamics parameters obtained from 3D doppler echocardiography. Our rationale was based on the extensive use of echo data used in the clinic, along with the rich literature showing the relationship between LVOT pressure gradients and DSS. In our results, we noted a significant increase LVOT mean and peak gradient between recurrent and non-recurrent DSS patients (**Figure 1**). This data was consistent with earlier published work^14^ and standard of care in the clinic, where an increase in LVOT mean and peak gradients are used to assess the condition of DSS patients. While LVOT mean and peak gradient continue to be used to evaluate and assess DSS patients, they do not provide any mechanistic insight or any insight into the pathways governing the progression of DSS. Further, LVOT mean and peak gradient cannot be used a predictive tool to assess the risk of re-occurrence of the fibrotic membrane in DSS patients. To overcome these limitations, we next sought to use differential changes in gene and protein expression from resected membranes from DSS patients.

Next, we sought to make use of data from RNA-sequencing to analysis resected fibrotic membranes from DSS patients (**Figures 2, 3 and 4**). Our data is presented in two formats, comparing patient samples at the time of first surgery, recurrent vs non-recurrent (**Figure 2 and 4**) and patient samples at the time of first and second surgery for recurrent DSS patients (**Figure 3 and Figure 4**). Analysis of the volcano plot for differential gene expression for DSS patients at the time of first surgery (**Figure 2A, Table S-1, S-2, Table 1**) identified multiple genes that are known to regulate ECM production and cell-matrix interactions. Over 20 such genes were identified, with notable examples including PKD2L1 (modulates cell matrix interactions), ACTR6 (cell structure and integrity) and PRKD1 (cell shape and adhesion). PCA analysis (**Figure 2B**) showed clustering and clear separation of the two groups, with the non-recurrent samples (blue) separating from the recurrent samples (red), suggesting differential molecular signatures between these two groups. The heatmap (**Figure 3C**) shows large red and blue areas, demonstrating significant changes in the differential gene expression between recurrent and nonrecurrent DSS patients at the time of first surgery.

There were changes in the gene profile of DSS patients between the first and second surgery for aggressive phenotype (**Figure 3 and 4**). The volcano (**Figure 3A**) shows the differential changes in all genes, also presented in Table S-3 and the 100 most significant genes presented in Table S-4. Relevant to our study hypothesis, the list of differential regulated genes related to ECM remodeling and turnover are presented in **Table 2** with NRAP (actin function), DSP (intermediate filament), CDH19 (cell-cell interaction) and CTHRC1 being important examples. The PCA analysis (**Figure 3B**) shows close clustering of samples from the two groups, suggesting lower differential gene expression between DSS patients at the time of surgery 1 vs surgery 2. The heatmap (**Figure 3C**) shows a similar result, which very little changes in the differential expression of genes between the two study groups.

Analysis of Hallmark and KEGG pathways (**Figure 4A, Table S-5, S-6**) identified epithelial to mesenchymal transition is a key modulator of DSS phenotype, for both groups (aggressive surgery 1 vs non-aggressive and aggressive surgery 2 vs 1). In addition, many pathways involved in ECM remodeling and function were also shown to be differentiated regulated based on Hallmark pathways, including TGF-beta signaling and protein secretion (**Figure 4A**). Similar results were obtained based on analysis of KEGG pathways (**Figure 4B**) for both study groups and significant changes in pathways regulating adherens junction, tight junction, focal adhesion and protein export being important examples. These results served as further validation for the critical role of ECM formation and turnaround is a key modulator of aggressive DSS phenotype. One interesting observation noted was the lower number of genes that are differentially expressed in aggressive DSS patients at the time of second surgery relative to first, based on gene ontology analysis and reactome pathways (**Figure 4C, Table S-7, S-8**). This suggest that many of the molecular changes in DSS patients, have already manifested by the time of first surgery, with little change between first and second surgery.

The volcano plots based on mass spec data is presented in **Figure 5**, **Table S-9-12**, **Table 3 and 4**. Collectively, this data serves to illustrate the role of ECM remodeling for the onset and progression of DSS. While the list of proteins that were differential regulated is extensive, important modulators were shown to be involved in actin remodeling, cytoskeleton organization, and cellular adhesion to underlying ECM. Gene ontology and pathway analysis (**Figure 6**, **Table S13-16**, **Table 5-8**) identified over 30 pathways related to ECM remodeling that are differentially regulated in DSS patients, with multiple aspects of actin formation, polymerization and filament organization differentially regulated (**Figure 6**). In-depth analysis of one specific pathway, related to the regulation of actin cytoskeleton (**Figure 7**), shows the critical points at which specific peptides are differentially enriched, providing insight into the molecular mechanisms. Collectively, the mass spec data provided insight into the role of ECM remodeling and organization on the onset and progression of DSS.

**Figure 6.**
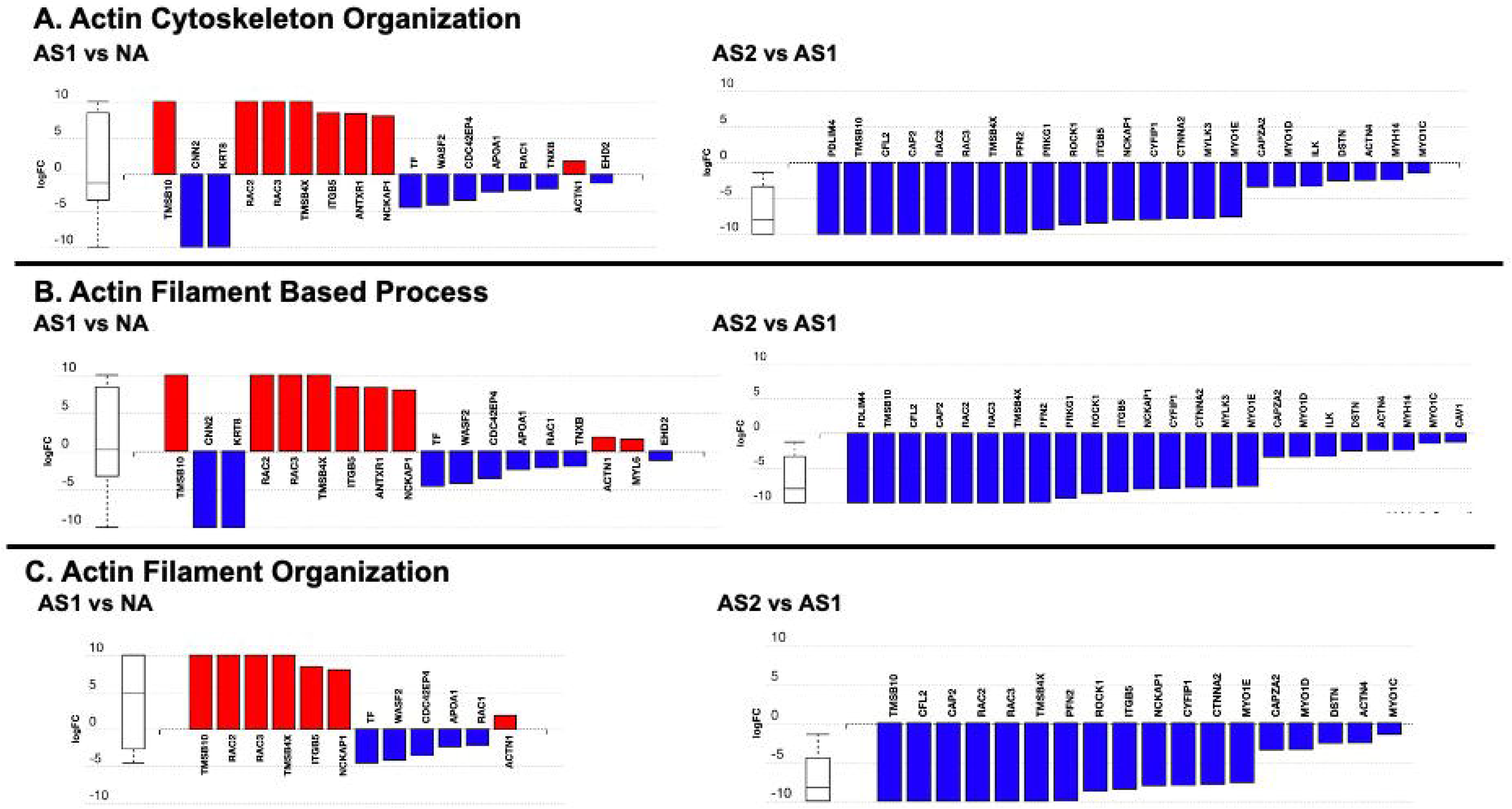
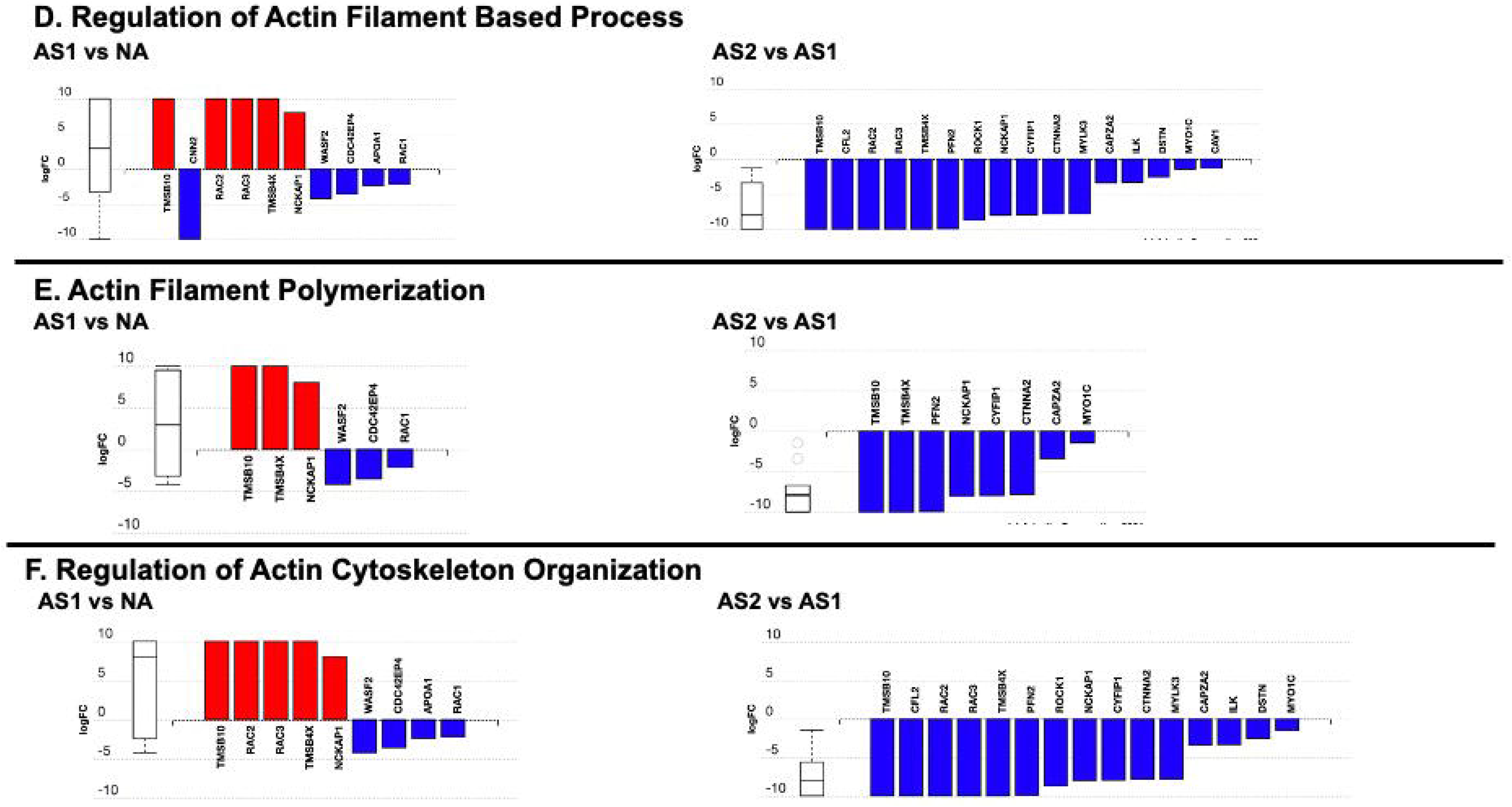
Gene Ontology Analysis Based on Proteomics Data. Differential regulations of individual peptides along six specific pathways related to actin assembly and remodeling are shown here. For each of the six pathways, the bar graph on the left shows differential expression at the time of first surgery, comparing AS1 vs NA, while the bar graph on the right compares AS2 vs AS1. **(A) Actin Cytoskeleton Organization**. **(B) Actin Filament Based Process**. **(C) Actin Filament Organization**. **(D) Regulation of Actin Filament Based Process**. **(E) Actin Filament Polymerization**. **(F) Regulation of Actin Cytoskeleton Organization**.

**Figure 7.**
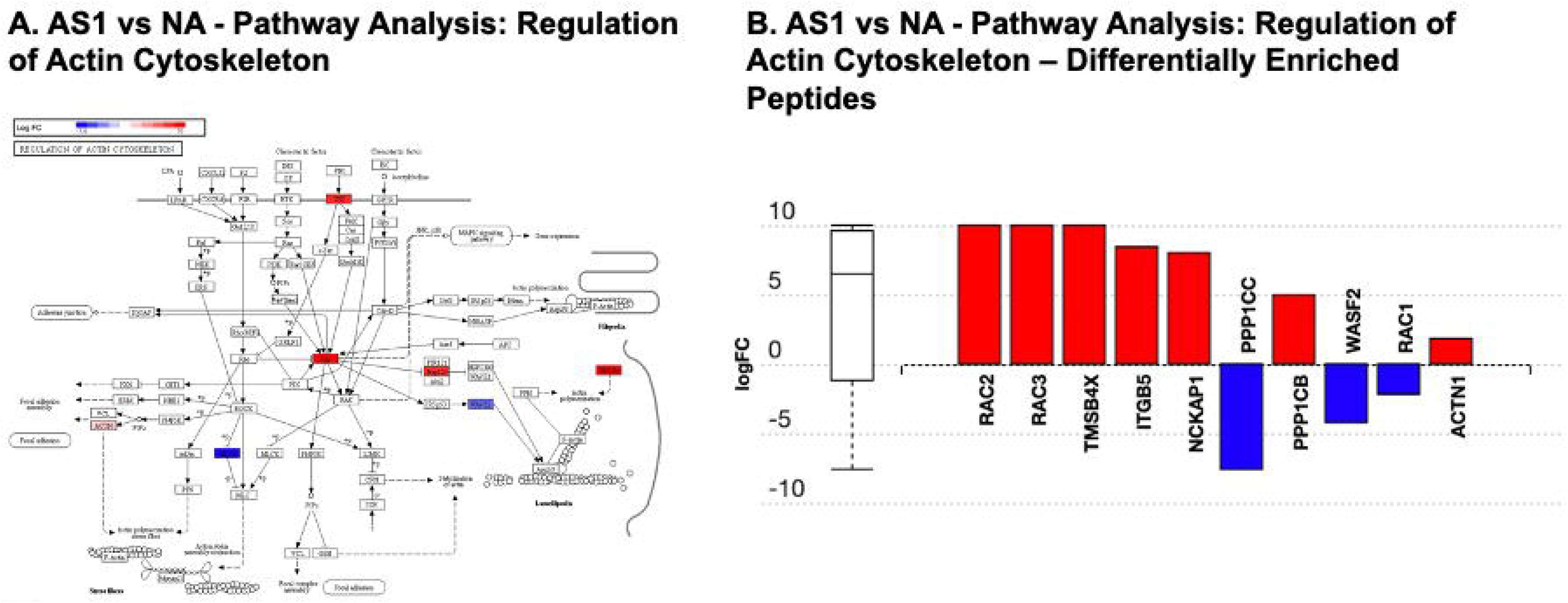
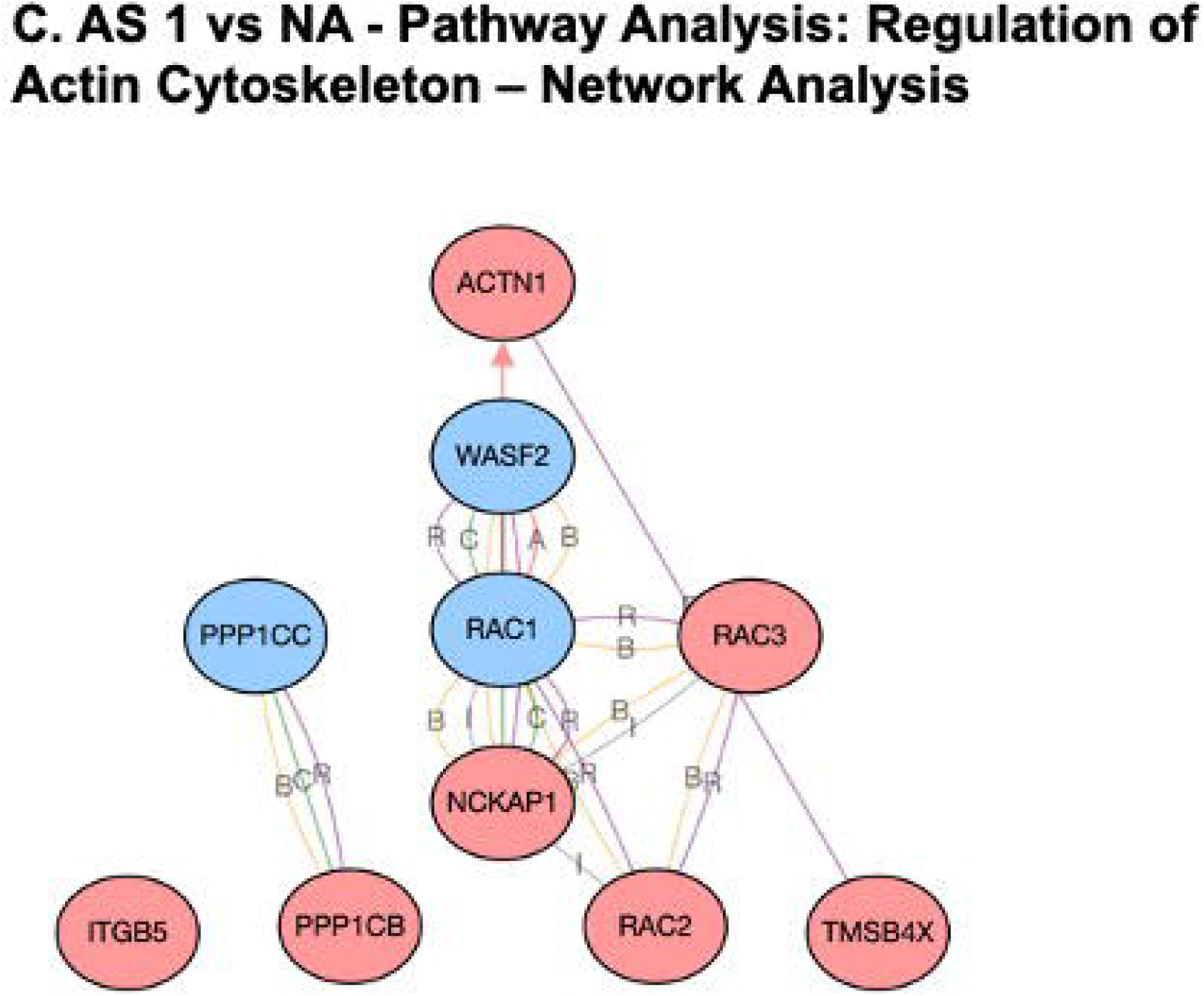
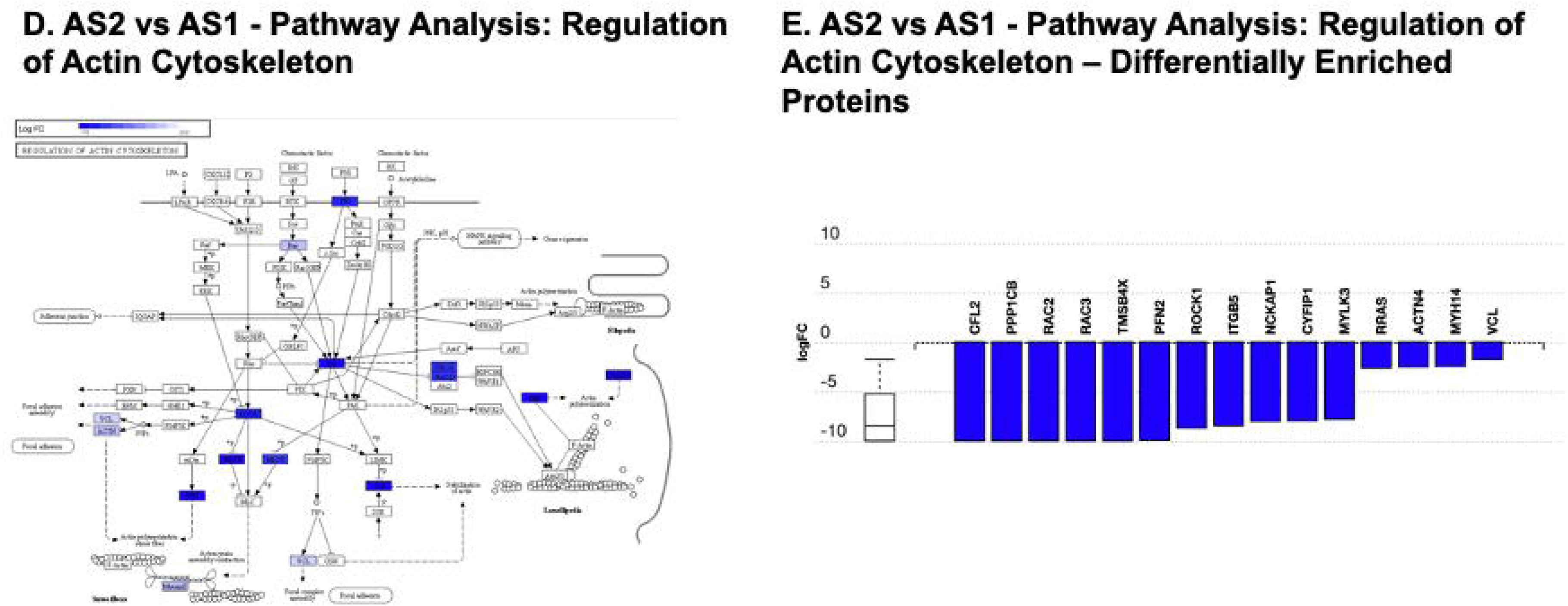
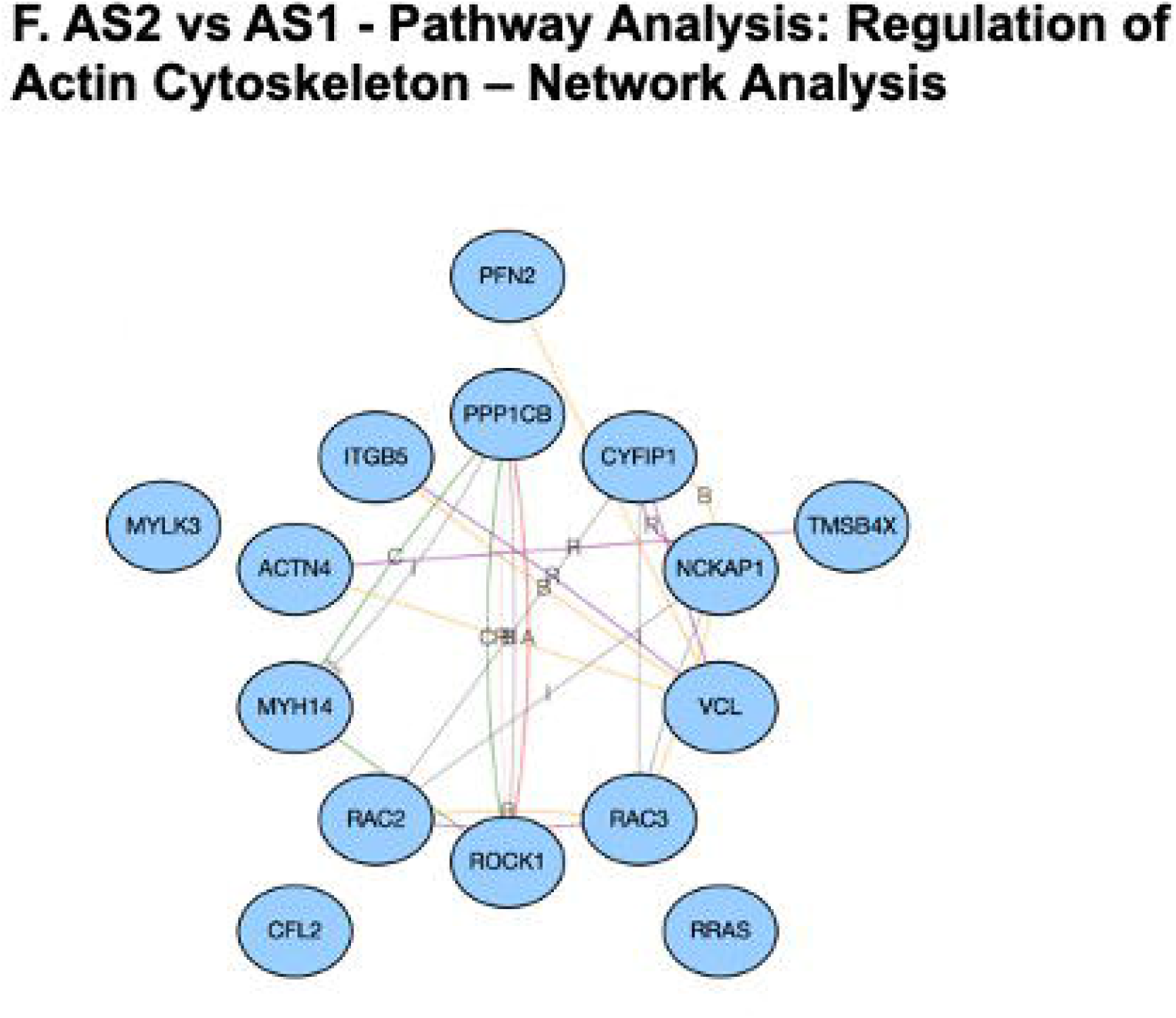
Regulation of Actin Cytoskeleton Based on Pathway Analysis for Proteomics Data. **(A-C)** Pathway Analysis, Differentially Enriched Peptides and Network Diagram for AS1 vs NA and **(D-F)** For AS2 vs AS1.

The results of our study support the role of ECM remodeling as a key modulator for the onset and progression of DSS. The ability to use molecular markers to predict the reoccurrence of DSS will provide a valuable tool for the care and management of DSS patients. This study is an important step in this direction. While this study demonstrates the role of ECM remodeling, further validation studies are needed to conclusively prove this prior to be using used a diagnostic tool in the clinic. Our echocardiography data presented in **Figure 1** shows an increase in LVOT peak and mean gradient in our DSS patient population, serving to validate this with published literature. Our genomic and proteomic data (**Figures 2-7**), show the role of ECM remodeling in DSS onset and progression, both at the gene (**Figure 2-4**) and protein (**Figures 5-7**) level.

## Supplemental Data

### List of Supplemental Tables

**Table S-1 – List of All Differentially Regulated Genes for AS1 vs NA at the time of First**

**Surgery.**

**Table S-2 – List of Top 100 Differentially Regulated Genes for AS1 vs NA Based on Significance.**

**Table S-3 – List of All Differentially Regulated Genes for AS2 vs AS1 for DSS Patients with an Aggressive Phenotype.**

**Table S-4 – List of Top 100 Differentially Regulated Genes for AS2 vs AS1 Based on Significance.**

**Table S-5 – Hallmark Pathway Analysis.**

**Table S-6 – KEGG Pathway Analysis.**

**Table S-7 – GOBP Pathway Analysis.**

**Table S-8 – Reactome Pathway Analysis.**

**Table S-9 – List of Differentially Regulated Peptides for AS1 vs NA Based on Proteomics Analysis.**

**Table S-10 – List of Top 100 Differentially Regulated Peptides for AS1 vs NA Based on Significance.**

**Table S-11 – List of Differentially Regulated Peptides for AS2 vs AS1 Based on Proteomics Analysis.**

**Table S-12 – List of Top 100 Differentially Regulated Peptides for AS2 vs AS1 Based on Significant.**

**Table S-13 - List of Differentially Regulated Pathways Based on Gene Ontology Analysis for AS1 vs NA.**

**Table S-14 – List of Differentially Regulated Pathways Based on Gene Ontology Analysis for AS2 vs AS1.**

**Table S-15 – List of Differentially Regulated Pathways Based on Pathway Analysis for AS1 vs NA.**

**Table S-16 – List of Differentially Regulated Pathways Based on Pathway Analysis for AS2 vs AS1.**

### List of Supplemental Figures

**Figure S-1 – Regulation of Adherens Junctions** – **(A-C) AS1 vs NA** – **(A)** Pathway Diagram, **(B)** Differential Enrichment of Peptides and **(C)** Network Diagram. **(D-F) AS2 vs AS1** – **(D)** Pathway Diagram, **(E)** Differential Enrichment of Peptides and **(F)** Network Diagram.

**Figure S-2 – Regulation of Focal Adhesions** – **(A-C) AS1 vs NA** – **(A)** Pathway Diagram, **(B)** Differential Enrichment of Peptides and **(C)** Network Diagram. **(D-F) AS2 vs AS1** – **(D)** Pathway Diagram, **(E)** Differential Enrichment of Peptides and **(F)** Network Diagram.

